# The nuclear transport receptor TNPO1 binds macrophage immunometabolism regulator MACIR via a PY-NLS motif

**DOI:** 10.1101/841999

**Authors:** Gavin McGauran, Emma Dorris, Razvan Borza, Niamh Morgan, Denis C. Shields, David Matallanas, Anthony G. Wilson, David J. O’Connell

## Abstract

Expression of the macrophage immunometabolism regulator gene (MACIR) is associated with severity of autoimmune disease pathology and the regulation of macrophage biology through unknown mechanisms. The 206 amino acid protein lacks homology to any characterized protein sequence and is a disordered protein according to structure prediction algorithms. Here we identify specific interactions of MACIR using a fragment complementation-based affinity pull down of cellular proteins prepared with a membrane solubilization buffer. Quantitative mass spectrometry showed enrichment of nuclear and mitochondrial proteins and of 63 significant interacting proteins, binding to the nuclear transport receptor TNPO1 and trafficking proteins UNC119 homolog A and B were validated by immunoprecipitation. Analysis of mutations in two candidate recognition motifs in the MACIR amino acid sequence confirmed TNPO1 binds via a PY-NLS motif (aa98-117). Characterizing nuclear MACIR activity in macrophage and fibroblasts is a priority with respect to developing strategies for treatment of autoimmune disease.

## Introduction

MACIR, previously named C5orf30, is a negative regulator of tissue damage and inflammation in rheumatoid arthritis (RA). It is highly expressed in the synovium of rheumatoid arthritis patients where it is predominately expressed by fibroblasts and macrophages (1). Reduced MACIR expression is an early event in the pathogenesis of RA and is negatively correlated with both Disease Activity Score for Rheumatoid Arthritis with ESR (DAS-ESR) and synovial Tumor Necrosis Factor (TNF) expression in early RA patients (2). Loss of expression contributes to the pathology of inflammatory arthritis *in vivo*, with inhibition of MACIR in the collagen-induced arthritis model leading to accentuated joint inflammation and tissue damage, and an increased inflammatory phenotype in a zebrafish model of wound healing (1,2). MACIR regulates macrophage cell phenotype, with increased MACIR expression found in M2-like anti-inflammatory macrophages. Its activity is regulated by multiple mechanisms in response to immune stimulants that include transcriptional and post-translational modifications and differential protein turn over (2). Inhibition of MACIR expression reduces wound healing/repair–associated functions of macrophages, reduces signaling required for resolution of inflammation, and decreases secretion of anti-inflammatory mediators (2). The ability of MACIR to negatively regulate inflammation is due, at least in part, to its role in regulating macrophage immunometabolism. In addition, recent evidence from the Human Pathology Atlas indicates that MACIR may be an unfavourable prognostic marker in liver and endometrial cancer. A CCNH: MACIR fusion was identified as a recurrent fusion gene across multiple cancer types (3–5). Thus, understanding more about the function of MACIR has wide ranging implications for health and disease.

MACIR is found exclusively in vertebrates and is tightly conserved between vertebrate species. However sequence analysis fails to identify homology to another vertebrate gene or protein. Modelling of protein secondary structure indicates that the protein is disordered without significant α-helical or β-sheet content preventing informative structure function analysis. Identifying mechanisms behind the function of this protein in the cell can be approached without this information through the identification of protein-protein interaction networks, their subcellular localization and the characterization of specific motifs and amino acids in the MACIR protein mediating these interactions.

In this study we have employed a membrane solubilization buffer (MSB) to enrich proteins from all cellular compartments from transfected HEK293T cells prior to pull down of affinity tagged MACIR and quantitative mass spectrometry analysis. We have previously developed a calcium-dependent fragment complementation-based affinity pull down system that used the highly specific recognition between two peptide fragments, EF1 & EF2, of a small calcium binding protein to pull down overexpressed kinase proteins from HEK293T cells and 7 transmembrane proteins from *Escherichia coli* membranes using MSB. This buffer contains a high concentration of glycerol and a cocktail of detergents to disrupt cellular and intracellular membranes and stabilise membrane proteins in detergent micelles (6,7). Protein complexes are eluted from the affinity matrix through calcium chelation and dissociation of the EF peptides prior to digestion and MS analysis, improving the signal to noise ratio for the identification of significant interactions. We then validate specific MACIR interactions with immunoprecipitation and use mutational analysis to characterize interacting motifs.

## Results & Discussion

### Affinity pull down of EF1-MACIR in RIPA & MSB

To determine the optimum conditions for pull down of MACIR interactors we performed western blot analysis of EF2 agarose pull downs of EF1-MACIR from normalised total protein lysates prepared with two cell lysis buffer formulations. MSB enriched more EF1-MACIR protein for pull down than RIPA buffer and the total amount of pulled down protein was also enhanced (Figure 1A i & ii). The empty vector control lysate and pull down for MSB is shown for comparison (Figure 1A iii). For mass spectrometry analysis proteins were eluted from the agarose prior to preparing peptides with tryptic digestion, followed by label-free quantitative mass spectrometry. Significant interactors were identified by dividing the average LFQ intensities found in the EF1-tagged MACIR samples by those identified in the EF1-tagged empty vector control. An LFQ intensity ratio greater than 2 with a p-value of less than 0.05 following t-test analysis was considered significant (8,9,10). A total of 9 proteins were identified as significant interactors in RIPA buffer pull downs, while these 9 proteins and 54 others were identified in MSB prepared lysates (Figure 1B, Table S1). RIPA buffer is commonly used to solubilize cells for immunoprecipitation and affinity pull down based mass spectrometry studies but it is clear from our data that a number of significant interactions with membrane bound proteins e.g., the mitochondrial ATP synthase subunits ATP5C1, ATP5L and ATP5A1, and proteins from membrane bound organelles e.g., nuclear PRKDC, would not have been identified without MSB cell lysis. The top 20 interactors identified with this approach, ranked by LFQ ratio, showed that in addition to the cytosolic proteins UNC119B and 14-3-3 proteins (YWAH, YWHAE and YWAHG) identified in both buffer conditions, the majority of interacting proteins were from nuclear and mitochondrial compartments identified in the MSB samples (Figure 1C).

**Figure 1.**
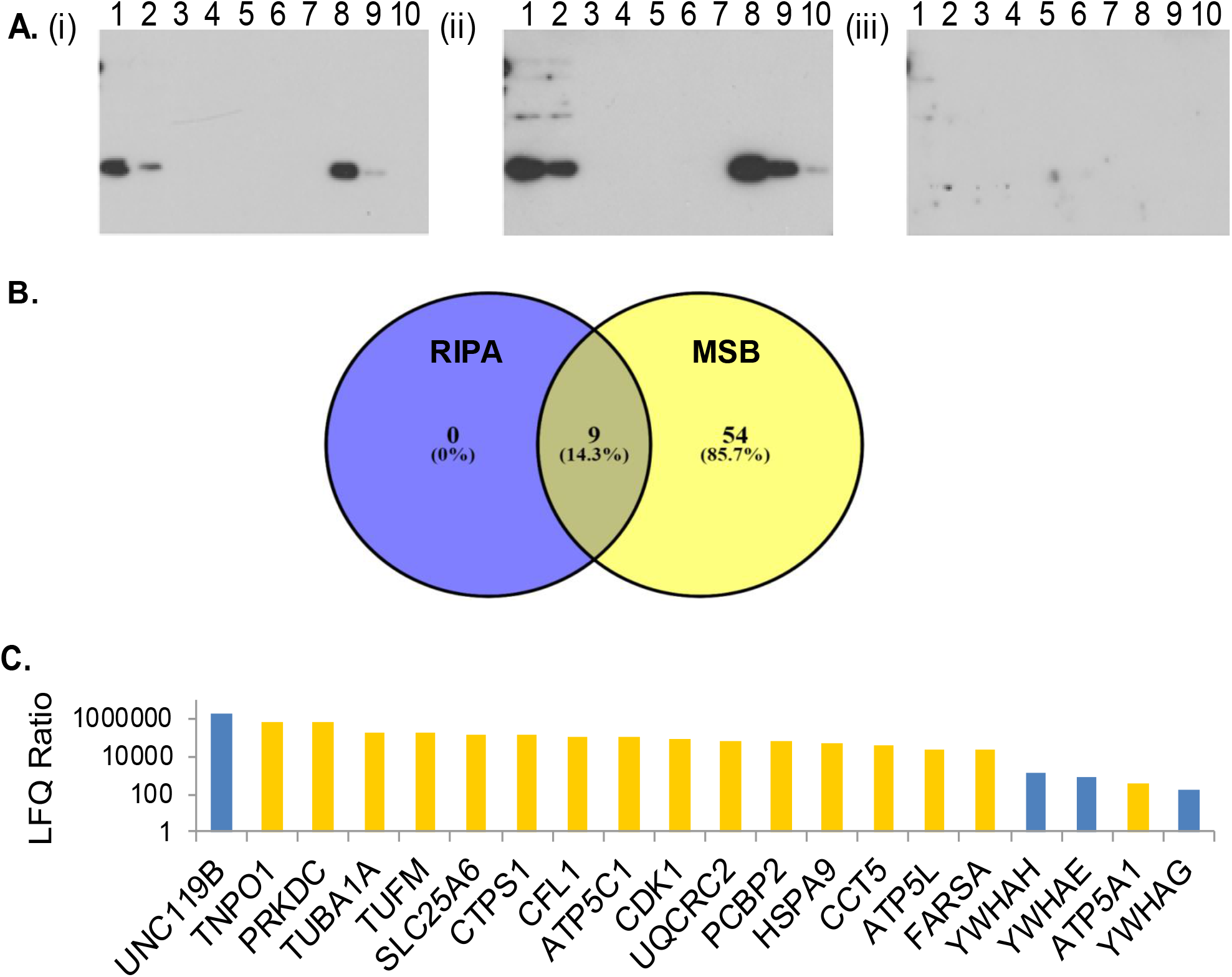
Proteomic analysis of MACIR binding interactions. (A) EF2 agarose pull down and elution of (i) EF1-tagged MACIR with RIPA buffer (ii) EF1-tagged MACIR with MSB (iii) EF1-tagged empty vector control with MSB (1:Input, 2:Unbound, 3-7:washes, 8-10:Eluates), probed with anti-EF1 polyclonal antibody. (B) Venn diagram of significant interactors identified with mass spectrometry in RIPA and MSB samples. (C) Top 20 MACIR interactors ranked by LFQ intensity ratio with proteins identified in MSB alone identified in yellow and in both buffers in blue.

14-3-3 protein isoforms are adapter proteins implicated in regulating a large number of general and specialized signaling pathways usually by recognition of a phosphoserine or phosphothreonine motif. They are likely to interact with MACIR via phosphoserine residues at positions 38 and 49 that are both confirmed in phosphopeptide mass spectrometry analysis (11) and are both predicted to be within 14-3-3 binding motifs (12) and manually curated linear motif database motifs (13).

### Immunoprecipitation validation of TNPO1 & UNC119 interactions with FLAG-MACIR

To confirm the specificity of the interactions identified with the mass spectrometry-based approach we next immunoprecipitated FLAG-tagged MACIR with anti-FLAG M2 sepharose beads and detected the co-precipitation of TNPO1 (Figure 2A i), and UNC119A and B (Fig 2A ii) from transfected HEK293T cell lysates. Anti-MACIR and anti-FLAG antibody controls confirmed specificity (Fig 2A ii) of the method. The interaction with UNC119 proteins confirms a previously reported interaction in a proteomics dataset in HEK293T cells (14). In the cell UNC119B has been shown to be required for localization of myristoylated proteins (15), while UNC119A cargo localization has been shown to regulate T-cell and macrophage signaling activities (16). TNPO1 is a transport receptor that transports substrates from the cytoplasm to the nucleus through nuclear pore complexes by recognizing nuclear localization signals (NLSs) and identification of this interaction raises the intriguing question of what is the function of MACIR in the nucleus. To confirm that it does localize to the nucleus in HEK293T cells we studied endogenous MACIR distribution with immunocytochemistry and confocal microscopy. Imaging of anti-MACIR stained HEK293T cells confirmed that MACIR is localized to the nucleus as well as the cytoplasm (Figure 2B).

**Figure 2.**
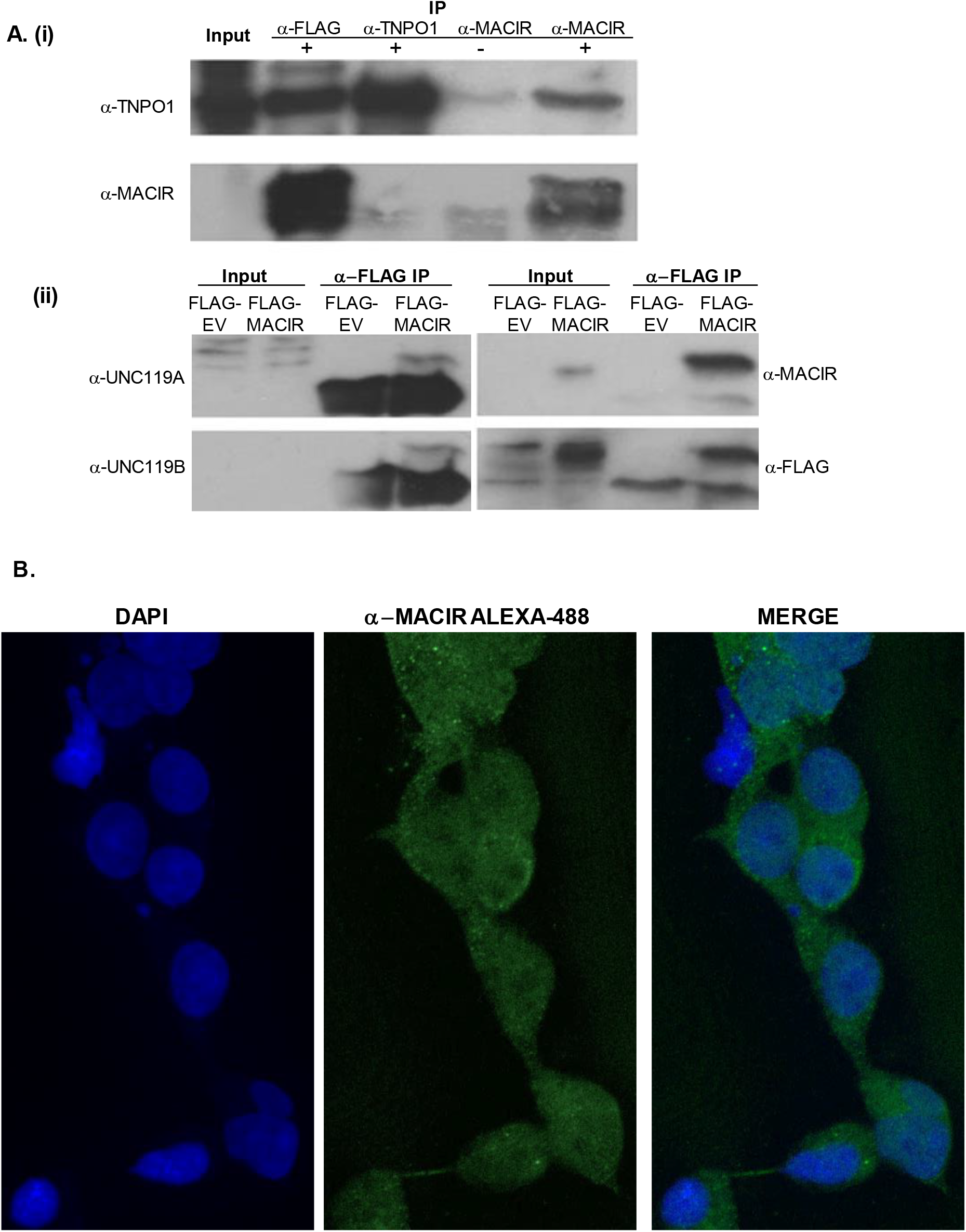
Validation of MACIR binding interactions and identification of nuclear localization of endogenous MACIR in HEK293T cells. (A) (i) Immunoprecipitation (IP) of FLAG-MACIR, TNPO1 and MACIR and western blot with anti-TNPO1 and anti-MACIR antibodies (ii) anti-FLAG IP of FLAG-MACIR and probing with anti-UNC119A and UNC119B antibodies and anti-FLAG antibody controls. (B) Confocal imaging of anti-MACIR staining in HEK293T cells at 60× magnification

**Figure 3.**
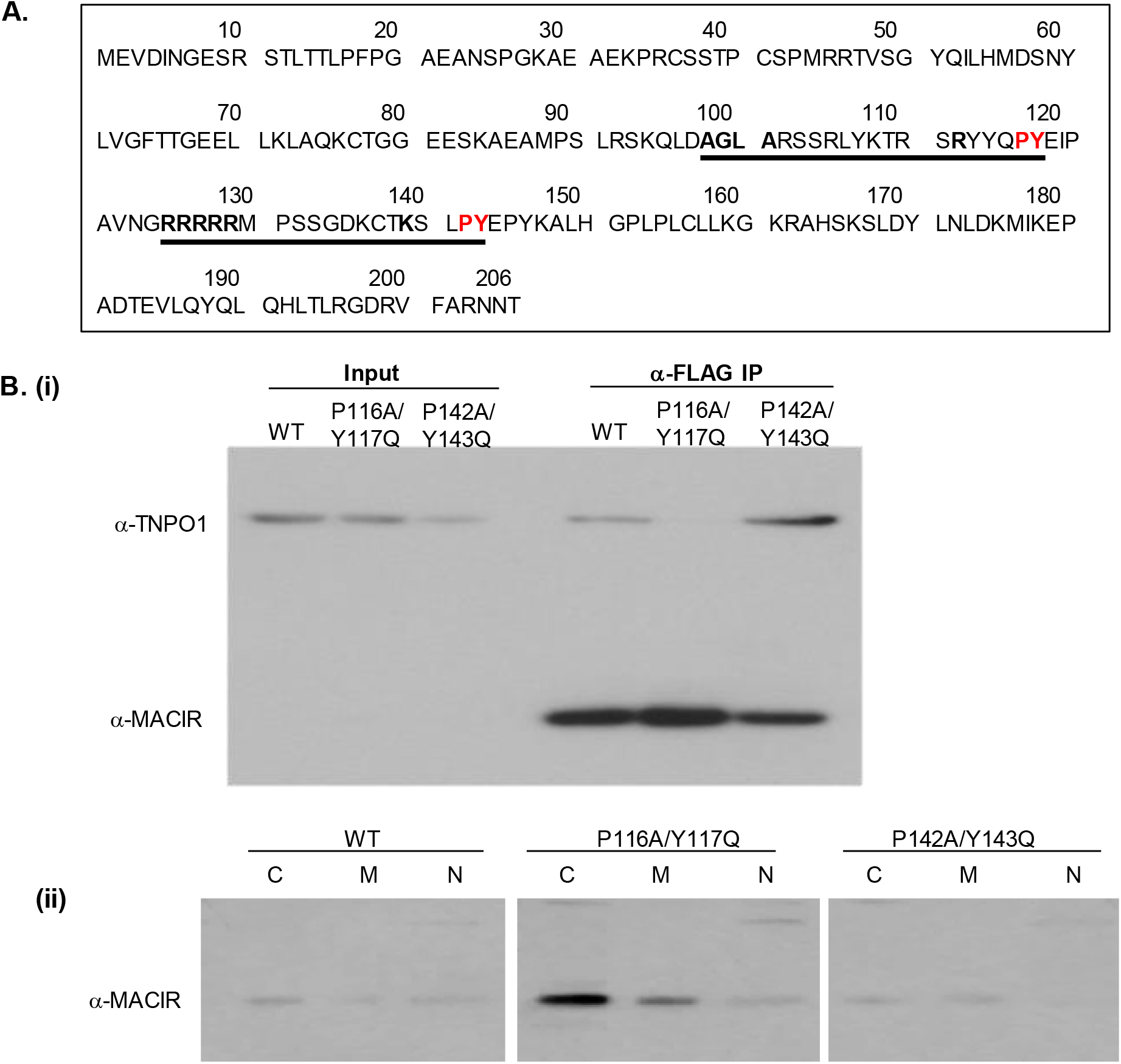
Mutational analysis of candidate TNPO1 recognition sites in MACIR. (A) Two candidate TNPO1 recognition sites studied underlined, with specific residues identified in the consensus sequence motif in bold and mutated PY residues shown in red. (B) (i) Anti-FLAG IP of FLAG-MACIR wt, P116A/117Q and P142A/Y143Q and probing with anti-TNPO1 and anti-MACIR (ii) Probing of subcellular fractions C (cytosol), M (membrane) and N (nuclear) with anti-MACIR.

TNPO1 recognizes cargo for import to the nucleus via PY-NLS motifs that are structurally disordered in free substrates, have overall basic character, and possess a central hydrophobic or basic motif followed by a C-terminal R/H/KX(2–5)PY consensus sequence (17). We analysed two candidate consensus TNPO1 binding sites in the primary sequence of MACIR (Fig 2A), that were previously described in other TNPO1 substrate proteins (17,18). We mutated proline/tyrosine (PY) residues to alanine/glutamine and analysed TNPO1 co-immunoprecipitation with FLAG-tagged MACIR P116A/Y117Q and MACIR P142A/Y143Q mutants. Our results show that TNPO1 binding to the MACIR P116/Y117Q mutant is lost while binding to P142A/Y143Q mutant is maintained, confirming that TNPO1 binds to MACIR via the PY-NLS recognition motif at amino acid positions 98-117 (Fig 2B i). Finally, we probed the subcellular fractions from these transfected cells with anti-MACIR and show a dysregulation of the subcellular localization of the P116A/Y117Q mutant with cytosolic accumulation of the mutant protein (Fig 2B ii).

### Conclusion

Our previous research has highlighted the potential for pro-resolution signaling in macrophages as a novel therapeutic target. Therapeutic agents that regulate localization of MACIR to particular sub-compartments may in the future yield pro-resolution treatments in inflammatory diseases such as rheumatoid arthritis. Identification of MACIR in the nucleus and characterization of a specific interaction with the nuclear receptor protein TNPO1 in this study highlights a promising avenue of future investigation in macrophage and fibroblast cells.

## Materials and Methods

### Chemicals and reagents

DNA PCR primers and construct sequencing were from MWG Eurofins Genomics (eurofinsgenomics.com). Restriction enzymes were from New England Biolabs (neb.com). EF2 agarose resin was from BlackLabBio (blacklabbio.com). Anti-MACIR (anti-C5orf30, HPA043434) polyclonal antibody, anti-UNC119A monoclonal antibody (SAB1404669), anti-FLAG monoclonal antibody (A8592) and anti-FLAG agarose were from Sigma (sigmaaldrich.com). Anti-EF1 polyclonal antibody (sc-28532) and Protein G agarose were from Santa Cruz (scbt.com). Anti-UNC119B polyclonal antibody (PA5-24504) was from ThermoFisher (thermofisher.com). Anti-TNPO1 monoclonal antibody was from AbCam (abcam.com). Cholesteryl hemisuccinate Tris salt (CHS) and detergents 3-[(3-cholamidopropyl) dimethylammonio]-1-propanesulfonate (CHAPS) and n-dodecyl-β-D-maltoside (DDM) were from Anatrace (anatrace.com).

### Cloning

EF1-tagged genes EF1-N-MACIR and EF1-C-MACIR were prepared by PCR. The forward primer: 5’-GGATCCGAAGTCGATATTAATG-3’ included a BamHI restriction site. The reverse primer: 5’-AAGCTTTGTATTATTCCTAGC-3’ included a HindIII restriction site for cloning into the expression vectors pEF1N-CMV and pEF1C-CMV. FLAG-tagged MACIR wt, P116A/Y117Q and P142A/Y143Q mutants with a C terminal FLAG tag were synthesized by Genscript (genscript.com).

### Cell culture and DNA transfection

HEK293T cells were seeded at 800,000 cells per 10 cm^3^ dish with Dulbecco’s Modified Eagle Media containing 10% foetal bovine serum (FBS) supplemented with penicillin and streptomycin. Cells were grown to 60-80% confluency prior to transfection. Transfection was performed as per manufacturer’s protocol (mirusbio.com) using 5 μg DNA and 2.4μl TransIT-2020/ μg of plasmid DNA made up in 1ml serum-free media. The transfection reagent:DNA complex mixture was incubated for 20 minutes at room temperature. Samples were then added drop-wise to pre-warmed media containing 10% FBS without penicillin and streptomycin and agitated. Cells were incubated at 37°C for 24 hours.

### Harvesting and lysis of cells

Cells were washed twice with PBS. Cells were harvested in 750 μl of RIPA (20 mM Tris-HCl, 150 mM NaCl, 1% Deoxycholic acid, 1% NP-40, 2 mM CaCl_2_, phosphatase and protease inhibitors) or MSB (50 mM Tris-HCl, 200 mM NaCl, 0.5% CHAPS, 0.1% CHS, 1% DDM, 30% Glycerol, 2 mM CaCl_2_, pH 7.5 and phosphatase and protease inhibitors) per 10 cm^3^ dish using a cell scraper. Cell pellets were mechanically disrupted via aspiration through a 21-gauge needle in the lysis buffer. Cells were centrifuged at 10,000 x g for 10 minutes at 4°C, retaining the supernatant. The total protein concentration was determined using bicinchoninic acid (BCA) assay (Pierce) and concentrations normalised for samples used in subsequent analyses.

### Affinity pull-down of EF1-MACIR from transfected lysates

EF2 agarose and unconjugated control beads were collected by centrifugation at 1,000 × g for 1 minute at 4°C. Beads were washed five times in 1 ml lysis buffer and 1.5 mg of lysate samples were precleared by incubation at 4°C for 1 hour using non-conjugated beads. The agarose beads were collected by centrifugation at 5,000 × g at 4°C for 1 minute and the supernatant was retained. The precleared lysate was incubated overnight at 4°C with EF2 agarose beads or non-conjugated beads as a negative control. Following incubation, samples were collected by centrifugation at 5,000 × g for 1 minute at 4°C. Samples were washed five times in lysis buffer. Following wash procedure, EF1-MACIR was eluted from the EF2-agarose beads using elution buffer (RIPA or MSB + 10 mM EDTA) by resuspension and allowing samples to incubate for 5 minutes at room temperature. Beads were collected by centrifugation at 5,000 × g for 1 minute and the supernatant was collected. Elution was repeated 5 times. Samples were stored at -80°C prior to further analysis.

### Mass Spectrometry Sample preparation

Samples were mixed with 400 μl of 8 M urea in 0.1 M Tris-HCl, pH 8.9 using a clean eppendorf. Samples were concentrated in Vivacon spin ultracentrifugation units at 14,000 × g for 15 minutes at 20°C for 15 minutes. 200 μl of 8 M urea in 0.1 M Tris-HCl, pH 8.9 was added to the filter unit and spun for 15 minutes at 20°C. The flow through was discarded and 100 μl of 0.05 M iodoacetamide/ 8 M urea in 0.1 M Tris-HCl, pH 8.9 added to the filter units. The samples were mixed for 1 minute at 600 rpm in a thermo-mixer and incubated in darkness for 20 minutes. A further spin at 14,000 × g for 10 minutes at 20°C was performed and then two washes with 100 μl of 8 M urea in 0.1 M Tris-HCl, pH 8.9, and a further two washes with 100 μl of 0.05 M NH_4_HCO_3_. The filter units were transferred to new collection tube and 40 μl of 0.05 M NH_4_HCO_3_ with trypsin (enzyme to protein ratio 1:50) added to the filter units and mixed at 600 rpm in a thermo-mixer for 1 minute. Filters were incubated on a wet chamber overnight at 37°C, spun at 14,000 × g for 15 minutes at 20°C to collect digested peptides. The filter units were washed once with 40 μl of 0.05 M NH_4_HCO_3_ to make a total volume of 80 μl per sample. The samples were measured with a NanoDrop™2000 to determine peptide concentration. c18 ZipTips were first equilibrated by aspirating 10 μl wetting solution (50% acetonitrile in 0.1% trifluoroacetic acid) twice prior to collection at 10,000 rpm for 5 min. Protein from the tryptic digest supernatants was aspirated and dispensed up to 10 times and washing solution (0.1% trifluoroacetic acid in ddH_2_O) was aspirated and dispensed twice through the ZipTip. 10 μL of elution solution (50% acetonitrile in 0.1% trifluoroacetic acid) was dispensed into a clean tube followed by aspiration and dispensing of the sample through the ZipTip at least 3 times without the introduction of air. Samples were evaporated on a CentriVap Concentrator for 10-15 min, allowing for a small retentate volume (6 μL). The sample was finally resuspended in 20 μL of 0.1% formic acid and centrifuged for 5 min at 15,000 rpm. 16 μL of this sample was transferred to a clean mass spectrometry vial for analysis.

### Mass spectrometry

Mass spectrometry was performed using an Ultimate3000 RSLC system that was coupled to an Orbitrap Fusion Tribrid mass spectrometer (Thermo Fisher Scientific). Following tryptic digest, the peptides were loaded onto a nano-trap column (300 μm i.d × 5mm precolumn that was packed with Acclaim PepMap100 C18, 5 μm, 100 Å; Thermo Scientific) running at a flow rate of 30 μl/min in 0.1% trifluoroacetic acid made up in HPLC water. The peptides were eluted and separated on the analytical column (75 μm i.d. × 25 cm, Acclaim PepMap RSLC C18, 2μm, 100 Å; Thermo Scientific) after 3 minutes by a linear gradient from 2% to 30% of buffer B (80% acetonitrile and 0.08% formic acid in HPLC water) in buffer A (2% acetonitrile and 0.1% formic acid in HPLC water) using a flow rate of 300 nl/min over 150 minutes. The remaining peptides were eluted using a short gradient from 30% to 95% in buffer B for 10 minutes. The mass spectrometry parameters were as follows: for full mass spectrometry spectra, the scan range was 335-1500 with a resolution of 120,000 at m/z=200. MS/MS acquisition was performed using top speed mode with 3 seconds cycle time. Maximum injection time was 50 ms. The AGC target was set to 400,000, and the isolation window was 1 m/z. Positive Ions with charge states 2-7 were sequentially fragmented by higher energy collisional dissociation. The dynamic exclusion duration was set at 60 seconds and the lock mass option was activated and set to a background signal with a mass of 445.12002.

### Analysis of mass spectrometry data

MS data analysis was performed using MaxQuant (version 1.5.3.30). Trypsin was set to be the digesting enzyme with maximal 2 missed cleavages. Cysteine carbamidomethylation was set for fixed modifications, and oxidation of methionine and N-thermal acetylation were specified as variable modifications. The data was then analysed with the minimum ratio count of 2. The first search peptide was set to 20, the main search peptide tolerance to 5 ppm and the 2re-quantify2 option was selected. For protein and peptide identification the Human subset of the SwissProt database (Release 2015_12) was used and the contaminants were detected using the MaxQuant contaminant search. A minimum peptide number of 1 and a minimum of 6 amino acids was tolerated. Unique and razor peptides were used for quantification. The match between run option was enabled with a match time window of 0.7 minutes and an alignment window of 20 minutes. A statistical analysis of the cut off ratio and two-sample t-test was done to determine interactors.

### Immunoprecipitation

0.5-1.5 mg cell lysates were used for immunoprecipitation experiments. Protein G agarose beads (Santa Cruz Biotechnology) or FLAG beads (Sigma) were collected by centrifugation at 10,000 × g for 30 seconds and washed three times in lysis buffer. 30 μl of FLAG beads or 30 μl protein G agarose beads were used per 0.5-1.5 mg of sample. The appropriate amount of antibody and equivalent amount of rabbit or mouse IgG control antibody was used per sample as per instructed by the manufactures data sheet. Samples were incubated with rotation for 1-4 hours at 4°C. After incubation samples were washed with lysis buffer three times. The beads containing the protein of interest were resuspended in 2× Laemmli loading buffer and boiled for 5 minutes and analysed by western blotting.

### Immunocytochemistry

Cells (25,000 per well) were seeded into 8-well chamber slides. Cells were fixed in 4% paraformaldehyde solution and permeabilised with 0.5% Triton × solution, washed in PBS and incubated with primary antibody (Rabbit Anti-MACIR 1/200 dilution) overnight at 4°C with agitation. Cells were washed in PBS and incubated with secondary antibody (Alexa-488 Anti-Rabbit 1/600 dilution) for two hours at room temperature, washed with PBS and a #1.5 coverslip mounted using Fluroshield™ with DAPI mounting media. Confocal images were acquired on a FluoView FV1000 laser scanning microscope (Olympus, Tokyo, Japan). Images were acquired with a 60×/1.35 NA UPlanSApo oil immersion objective (Olympus, Tokyo, Japan) at a resolution of 1024 × 1024 pixels in sequential scanning model. Images were pseudo-coloured in ImageJ.

## Acknowledgements

We gratefully acknowledge the support of the Health Research Board & Arthritis Ireland grant MRCG-2016-15, the Irish Research Council award IRC: GOIPD/2017/1310 and Science Foundation Ireland award 16/RC/3998 (BEACON).

## Supplementary Information

**Table S1.**
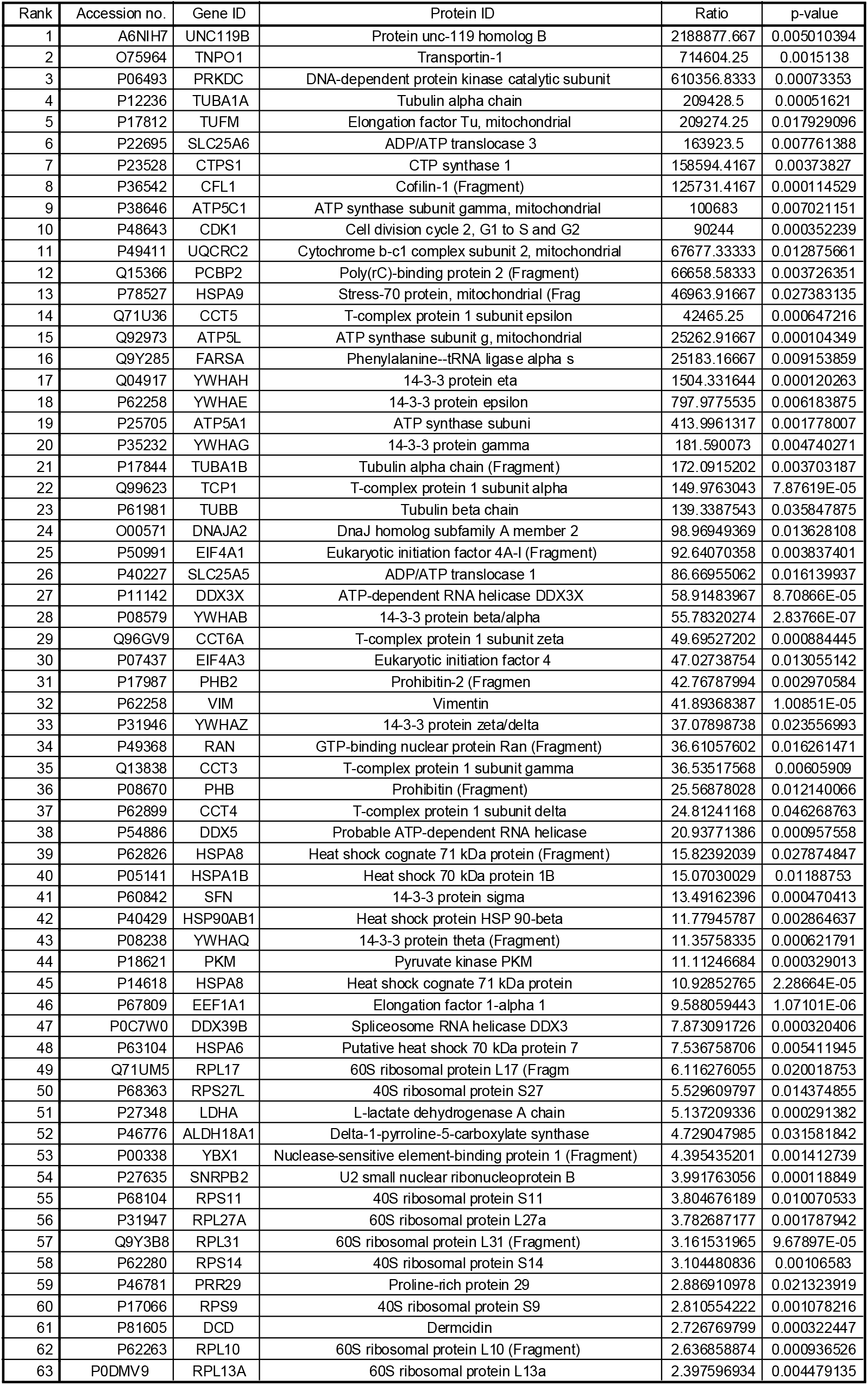
Statistically significant MACIR interacting proteins. Label free quantitative mass spectrometry data ranked following the application of an average intensity ratio EF1-MACIR/EF1 control cut-off above 2 and p-value of less than 0.05.

## References

1. Muthana M, Hawtree S, Wilshaw A, Linehan E, Roberts H, Khetan S, Adeleke G, Wright F, Akil M, Fearon U, Veale D, Ciani B, Wilson AG. C5orf30 is a negative regulator of tissue damage in rheumatoid arthritis. Proc Natl Acad Sci U S A. 2015 Sep 15;112(37):11618–23.

2. Dorris ER, Tazzyman SJ, Moylett J, Ramamoorthi N, Hackney J, Townsend M, Muthana M, Lewis MJ, Pitzalis C, Wilson AG. The Autoimmune Susceptibility Gene C5orf30 Regulates Macrophage-Mediated Resolution of Inflammation. J Immunol. 2019 Feb 15;202(4):1069–1078.

3. Uhlen M, Zhang C, Lee S, Sjöstedt E, Fagerberg L, Bidkhori G, Benfeitas R, Arif M, Liu Z, Edfors F, Sanli K, von Feilitzen K, Oksvold P, Lundberg E, Hober S, Nilsson P, Mattsson J, Schwenk JM, Brunnström H, Glimelius B, Sjöblom T, Edqvist PH, Djureinovic D, Micke P, Lindskog C, Mardinoglu A, Ponten F. A pathology atlas of the human cancer transcriptome. Science. 2017 Aug 18;357(6352).

4. Yu YP, Liu P, Nelson J, Hamilton RL, Bhargava R, Michalopoulos G, Chen Q, Zhang J, Ma D, Pennathur A, Luketich J, Nalesnik M, Tseng G, Luo JH. Identification of recurrent fusion genes across multiple cancer types. Sci Rep. 2019 Jan 31;9(1):1074.

5. Yu YP, Tsung A, Liu S, Nalesnick M, Geller D, Michalopoulos G, Luo JH. Detection of fusion transcripts in the serum samples of patients with hepatocellular carcinoma. Oncotarget. 2019 May 21;10(36):3352–3360.

6. Dunning CJ, McGauran G, Willén K, Gouras GK, O’Connell DJ, Linse S. Direct High Affinity Interaction between Aβ42 and GSK3α Stimulates Hyperphosphorylation of Tau. A New Molecular Link in Alzheimer’s Disease? ACS Chem Neurosci. 2016 Feb 17;7(2):161–70.

7. Mhurchú NN, Zoubak L, McGauran G, Linse S, Yeliseev A, O’Connell DJ. Simplifying G Protein-Coupled Receptor Isolation with a Calcium-Dependent Fragment Complementation Affinity System. Biochemistry. 2018 Jul 31;57(30):4383–4390.

8. Turriziani B, Garcia-Munoz A, Pilkington R, Raso C, Kolch W, von Kriegsheim A. On-beads digestion in conjunction with data-dependent mass spectrometry: a shortcut to quantitative and dynamic interaction proteomics. Biology (Basel). 2014 Apr 16;3(2):320–32.

9. Rodriguez J, Herrero A, Li S, Rauch N, Quintanilla A, Wynne K, Krstic A, Acosta JC, Taylor C, Schlisio S, von Kriegsheim A. PHD3 Regulates p53 Protein Stability by Hydroxylating Proline 359. Cell Rep. 2018 Jul 31;24(5):1316–1329.

10. Rodriguez J, von Kriegsheim A. Mass Spectrometry and Bioinformatic Analysis of Hydroxylation-Dependent Protein-Protein Interactions. Methods Mol Biol. 2018;1742:27–36.

11. Hornbeck PV, Kornhauser JM, Tkachev S, Zhang B, Skrzypek E, Murray B, Latham V, Sullivan M. PhosphoSitePlus: a comprehensive resource for investigating the structure and function of experimentally determined post-translational modifications in man and mouse. Nucleic Acids Res. 2012;40(Database issue):D261–70.

12. Madeira F, Tinti M, Murugesan G, Berrett E, Stafford M, Toth R, Cole C, MacKintosh C, Barton GJ. 14-3-3-Pred: improved methods to predict 14-3-3-binding phosphopeptides. Bioinformatics. 2015 15;31(14):2276–83.

13. Marc Gouw, Sushama Michael, Hugo Sámano-Sánchez, Manjeet Kumar, András Zeke, Benjamin Lang, Benoit Bely, Lucía B Chemes, Norman E Davey, Ziqi Deng, Francesca Diella, Clara-Marie Gürth, Ann-Kathrin Huber, Stefan Kleinsorg, Lara S Schlegel, Nicolás Palopoli, Kim V Roey, Brigitte Altenberg, Attila Reményi, Holger Dinkel, Toby J Gibson. The eukaryotic linear motif resource – 2018 update. Nucleic Acids Research, Volume 46, Issue D1, 4 January 2018, Pages D428–D434

14. Huttlin EL, Bruckner RJ, Paulo JA, Cannon JR, Ting L, Baltier K, Colby G, Gebreab F, Gygi MP, Parzen H, Szpyt J, Tam S, Zarraga G, Pontano-Vaites L, Swarup S, White AE, Schweppe DK, Rad R, Erickson BK, Obar RA, Guruharsha KG, Li K, Artavanis-Tsakonas S, Gygi SP, Harper JW. Architecture of the human interactome defines protein communities and disease networks. Nature 2017 May 25;545(7655):505–509.

15. Wright KJ, Baye LM, Olivier-Mason A, Mukhopadhyay S, Sang L, Kwong M, Wang W, Pretorius PR, Sheffield VC, Sengupta P, Slusarski DC, Jackson PK. An ARL3-UNC119-RP2 GTPase cycle targets myristoylated NPHP3 to the primary cilium. Genes Dev. 2011 25:2347–2360.

16. Gorska MM1, Goplen N, Liang Q, Alam R. Uncoordinated 119 preferentially induces Th2 differentiation and promotes the development of asthma. J Immunol. 2010 15;184(8):4488–96.

17. Lee BJ, Cansizoglu AE, Süel KE, Louis TH, Zhang Z, Chook YM. Rules for nuclear localization sequence recognition by karyopherin beta 2. Cell. 2006 Aug 11;126(3):543–58.

18. Imasaki T, Shimizu T, Hashimoto H, Hidaka Y, Kose S, Imamoto N, Yamada M, Sato M. Structural basis for substrate recognition and dissociation by human transportin 1. Mol Cell. 2007 Oct 12;28(1):57–67.

